# Antimicrobial susceptibility patterns of and biofilm formation by *Staphylococcus aureus* strains isolated from pediatric patients with atopic dermatitis

**DOI:** 10.1101/2025.03.28.645872

**Authors:** Romo-González Carolina, Aquino-Andrade Alejandra, Perez-Carranza Abril, Chaparro-Camacho Diana, Becerril-Osnaya Andrea, Garcia-Romero Maria Teresa

**Affiliations:** Experimental Bacteriology Laboratory, Instituto Nacional de Pediatria, Mexico City. México; Molecular Microbiology Laboratory, Instituto Nacional de Pediatria, Mexico City. México; Diagnostic Bacterology, Facultad de Estudios Superiores Cuautitlan, UNAM. Estado de México. México; Department of Dermatology, Instituto Nacional de Pediatria, Mexico City. México

## Abstract

**Background:** Atopic dermatitis (AD) is a chronic inflammatory skin disease characterized by barrier dysfunction and susceptibility to *Staphylococcus aureus* colonization. Biofilm formation alters antibiotic resistance and the immune response. This study examines the antimicrobial susceptibility patterns and biofilm formation of *S. aureus* isolates from pediatric AD patients.

**Methods:** A prospective longitudinal observational study was conducted with 136 *S. aureus* isolates collected from 26 pediatric patients with moderate-to-severe AD over 18 months. Isolates were obtained from both lesional and nonlesional skin and from nares. Antimicrobial susceptibility was evaluated using the disk diffusion method, whereas biofilm production was quantified using a crystal violet microtiter assay. Clinical characteristics, including AD severity, response to treatment, and the use of adjunctive dilute bleach baths, were analyzed for associations with *S. aureus* features.

**Results:** Of the isolates, 60.2% exhibited moderate-to-strong biofilm production, which was associated with severe AD at baseline (p=0.01), lack of clinical improvement (p=0.04), and persistent moderate-severe disease (p=0.01). The antimicrobial resistance rates for penicillin, gentamicin, clindamycin, and erythromycin exceeded 15%. Isolates from patients receiving dilute bleach baths presented greater resistance to ciprofloxacin (p<0.0001) and constitutive and inducible macrolide-lincosamide-streptogramin B resistance (MLSB). Inducible MLSB was associated with the *ermA* gene in 80% of the cases.

**Conclusions:** *S. aureus* biofilm formation correlates with disease severity and treatment resistance in pediatric AD patients. These findings highlight the need for culture-guided therapy and emphasize the importance of tailored strategies to manage *S. aureus* colonization and infection in AD patients.

**Importance:** In this longitudinal study of 136 *S. aureus* isolates from 26 pediatric patients with atopic dermatitis, we examined the antimicrobial susceptibility patterns and biofilm formation of *S. aureus*. We found 60.2% exhibited moderate-to-strong biofilm production, which was significantly associated with severe atopic dermatitis at baseline, lack of clinical improvement, and persistent moderate-severe disease. The antimicrobial resistance rates for penicillin, gentamicin, clindamycin, and erythromycin were suboptimal exceeding 15%. Isolates from patients receiving dilute bleach baths presented greater resistance to ciprofloxacin and constitutive and inducible MLSB resistance. Inducible MLSB resistance was associated with the *ermA* gene in 80% of the cases. These findings emphasize the importance of culture-guided therapy in skin and soft tissue infections associated to atopic dermatitis, the need for tailored strategies to manage *S. aureus* colonization and infection in AD patients, and of developing alternative treatments targeting biofilm and quorum-sensing mechanisms.

## Introduction

Atopic dermatitis (AD) is a chronic multifactorial inflammatory skin disease characterized by alterations in epidermal barrier function and an exaggerated innate immune response. It affects 20–30% of children and is one of the most frequent drivers of medical consultation (1, 2). Manifestations of AD include erythematous scaly patches that are intensely pruritic and itchy, leading to a vicious cycle of scratching that exacerbates the disease. Patients frequently exhibit colonization of superinfection by *Staphylococcus aureus*, and this pathogen plays a role in the physiopathology of the disease by producing superantigens, decreasing the expression of antimicrobial peptides and forming biofilms that impair the effects of targeted treatment (1, 2).

The antimicrobial susceptibility of *S. aureus* strains that cause disease or have been isolated from patients with AD has been reported to differ from that of *S. aureus* strains isolated from healthy controls. Since differentiating mild superinfection from disease flares may be challenging, antibiotics are frequently used as empiric treatment (3). Furthermore, resistance to antibiotics recommended as first-line therapy for skin infections may be suboptimal (4–9) and methicillin resistance and multidrug resistance are potentially associated with age, severity of AD and treatment (6, 10).

*S. aureus* produces biofilms, which are bacterial agglomerations that are attached to a surface and embedded in an extracellular matrix. The formation of biofilms successfully protects bacteria from environmental hazards, innate immune response-derived antimicrobial peptides (AMPs), antibiotics, and phagocytosis and contributes to the pathogenesis of chronic infections. Biofilms have been detected on the skin of AD patients and possibly contribute to the inflammation process (11–14).

In this study, we sought to analyze the antimicrobial susceptibility profiles and characterize the biofilm formation of *S. aureus* strains isolated longitudinally from children with AD, as well as identify differences associated with clinical characteristics.

## Materials and methods

This was a prospective longitudinal observational study performed at the National Institute of Pediatrics (NIP) (IRB approval number 073/2019). We obtained *S. aureus* isolates from a cohort of children with moderate–severe AD who came to the dermatology clinic from July 2017 to December 2018 and consented to participate. The severity of AD was quantified via the Severity Score for AD (SCORAD) (15). Superficial swabs were obtained from 7 body sites (nares, antecubital folds and popliteal folds) in up to 5 visits (one baseline and 4 follow-up). Cultures and colony isolation were performed at the Experimental Bacteriology Laboratory of the NIP. From each patient, we selected up to 3 colonies of one isolate obtained from a lesional site, one from a nonlesional site, and one from nares at baseline and at a follow-up visit (1 to 4 months after). Isolates that had been obtained from initial visits were given preference over those from the latest visit. For each isolate, we 1) performed antimicrobial susceptibility testing via the disc diffusion test, 2) analyzed *in vitro* biofilm production, and 3) performed detection of the *erm*A, *erm*B, *erm*C and *msr*A genes by PCR. Patients were categorized as those receiving standard treatment for their AD (medium- to high-potency topical steroids and/or calcineurin inhibitors) and those receiving adjunctive dilute bleach baths (0.006%).

### Sample collection and processing

Each sample was then streaked onto the surface of sheep blood agar and mannitol salt agar plates and incubated at 37°C for 24 h for purity verification and identification. After the incubation period, colonies were identified based on mannitol fermentation and colony morphology in addition to Gram staining, and the final identification was performed using conventional biochemical tests (e.g., catalase, coagulase, and DNase tests).

A total of 136 *S. aureus* isolates from patients with AD were collected from different sites. All the isolates were further tested for antimicrobial susceptibility and biofilm formation.

### Antimicrobial susceptibility testing

The antimicrobial susceptibility profiles and detection of inducible clindamycin resistance (ICR) test were performed following the guidelines of the Clinical and Laboratory Standards Institute (16, 17). A disk diffusion test was performed for penicillin (P), cefoxitin (FOX), erythromycin (E), clindamycin (CC), ciprofloxacin (CIP), tetracycline (TE), gentamicin (GM), trimethoprim-sulfamethoxazole (SXT), and linezolid (LIN) (Becton Dickinson, Franklin Lakes, Nueva Jersey, USA). The iMLSB was considered when the isolates were resistant to E or sensitive or intermediate to CC, with a positive ICR test; the MSB phenotype was considered when the ICR test was negative; and the constitutive phenotype (cMLSB) was considered if they were resistant to E and CC (18).

### Detection of erm*A*, erm*B*, erm*C* and msr*A* genes

DNA was obtained with the QIAmp DNA mini® kit (Qiagen, Hilden, North Rhine– Westphalia, Germany). The DNA was eluted and stored at −20°C until use. The genes *erm*A, *erm*B, *erm*C and *msr*A were amplified by PCR using previously reported primers and conditions (19, 20).

### Assay biofilm

Each isolate was grown on brain heart infusion (BHI) agar plates at 37°C for 24 h. Next, the cell density was adjusted to a concentration of 0.5 MacFarland scale (1.5 × 10^8^ CFU/mL) in 0.9% physiological saline solution. The strain controls for this test were the same as those used by Singh et al. (21)

Biofilm production was performed with BHI supplemented with 222.2 mM glucose, 116.9 mM saccharose and 1000 mM NaCl. Overnight cultures were enriched with BHI, a suspension equivalent to 0.5 MacFarland was diluted 1:100 in freshly prepared BHI. Aliquots (100 μL) were transferred to 96-well microtiter plates. Sterile broth without culture served as a control. The plates were incubated at 37°C for 48 h. After incubation, the contents of each well were gently decanted. The wells were washed 2–3 times with 200 μL of phosphate-buffered saline (PBS) to remove planktonic bacteria. Then, the plate was air dried at room temperature and stained with 0.5% crystal violet. The wells were subsequently washed with distilled water 5–6 times to remove excess stain. The stain adherent to the walls was dissolved in 100 μL of alcohol:acetone (80:20), and the optic density was read at 570 nm via an Epoch Microplate Absorbance Reader (Agilent, USA). The biofilms formed by the *S. aureus* isolates were classified as strong, moderate, weak, or none, as suggested by Singh et al. (21)

### Patientś clinical characteristics

We investigated differences in the antimicrobial susceptibility patterns and biofilm formation of *S. aureus* isolates according to the following categories based on the clinical characteristics of the patients:

1. Body sites from which isolates were obtained: affected skin, unaffected skin, or nares
2. Severity of AD at baseline: moderate or severe
3. Baseline visit or follow-up visit (once patients had received treatment)
4. Number of body sites colonized by *S. aureus* at baseline: ≤ 3 sites or more than 3 sites
5. Adjunctive treatment with or without dilute bleach baths
6. Response to treatment (decrease of at least 15 SCORAD points from baseline to final visit)
7. Persistent moderate-to-severe AD throughout all visits

### Statistical analysis

Descriptive statistics were used to summarize continuous variables with means and standard deviations and categorical variables with numbers and percentages. Differences between groups were analyzed via the chi-square test or Fisher’s exact test (SPSS Version 21.0). A p value of 0.05 was considered significant.

## Results

### Characteristics of the *S.* aureus isolates

We included 136 isolates of *S. aureus* from 26 patients (Table 1). Fifty-five (40.4%) of the isolates were obtained from patientś nares, 50 (36.7%) from affected skin, and 31 (22.7%) from unaffected skin. All patients had moderate or severe AD at baseline (SCORAD ≥25). At the baseline visit, all patients were prescribed treatment with emollients and moderate-potency topical steroids twice a day; 12 received adjunctive treatment with diluted bleach baths (concentration 0.006%) in addition to standard treatment, while 14 did not.

**Table 1.** Clinical characteristics of 26 patients from whom *S. aureus* was isolated (n = 136)

| Clinical characteristics | Number of isolates (%) |
| --- | --- |
| Site |  |
| Affected skin | 50 (36.7) |
| Unaffected skin | 31 (22.7) |
| Nares | 55 (40.4) |
| Severity of AD at baseline<sup>a</sup> |  |
| Moderate | 42 (58.3) |
| Severe | 30 (41.7) |
| Visit |  |
| Baseline | 72 (52.9) |
| Subsequent | 64 (47) |
| Body sites colonized by *S. aureus* at baseline<sup>a</sup> |  |
| ≤ 3 | 14 (19.4) |
| > 3 | 58 (80.6) |
| Adjunctive treatment with dilute bleach baths<sup>b</sup> |  |
| No | 41 (62.1) |
| Yes | 23 (37.9) |
| SCORAD decreased 15 or more points from baseline to 4th visit |  |
| No | 40 (29.4) |
| Yes | 96 (70) |
| Persistently moderate-severe AD |  |
| No | 78 (57.3) |
| Yes | 58 (42.7) |

Abbreviations: SCORAD: Severity score for AD; AD: atopic dermatitis.
<sup>a</sup> Only samples from the baseline visit were analyzed (n = 72).
<sup>b</sup> Only samples from subsequent visits were analyzed (n = 66).

Seventy-two isolates of *S. aureus* (52.9%) were obtained from the baseline visits. At the initial visit, 15 patients had >3 body sites colonized by *S. aureus*, and 58 isolates (80.5%) were obtained from these patients.

Sixty-four isolates (47%) were obtained from follow-up visits. Of these, 23 (35.9%) were isolated from 7 patients who received adjunctive treatment with diluted bleach baths, and 41 (64%) were isolated from 9 patients who did not.

Most patients (18, 69.2%) responded well to treatment, and their disease severity decreased 15 or more SCORAD points from baseline to the end of the study; 96 isolates (70.5%) were derived from these individuals. Five patients (19.2%) had persistent moderate–severe AD throughout the study period, accounting for 58 isolates (42.6%).

### Antimicrobial susceptibility patterns and biofilm production

In general, isolates of *S. aureus* had suboptimal rates of resistance (>15%) to P (103, 75.7%), GM (49, 36%), CC (34, 25%), and E (34, 25%). All 136 isolates were methicillin sensitive.

The MLSB phenotype was documented in 58 (23.8%) of the isolates: 14 (10.2%) had cMLSB, 20 (14.7%) had iMLSB, and 1 (0.7%) had the MSB phenotype (Table 2). The *erm*A gene was detected in 16 isolates (80%) with the iMLSB phenotype, one of which was associated with the *msr*A gene; *erm*B and *erm*C were not detected.

**Table 2.** Patterns of susceptibility/resistance to the tested antibiotics in all the *S. aureus* isolates (n = 136)

| Variables | LZ | CC | GM | CIP | P | E | TE | SXT | FOX |
| --- | --- | --- | --- | --- | --- | --- | --- | --- | --- |
| Resistant, n (%) | 0 | 34 (25) | 87 (64) | 1 (0.7) | 103 (75.7) | 34 (25) | 3 (2.2) | 1 (0.7) | 0 |
| Intermediate, n (%) | – | – | – | 21 (15.4) | – | 1 (0.7) | 9 (6.6) | 0 | – |
| Sensitive, n (%) | 136 (100) | 102 (75) | 49 (36) | 114 (83.8) | 33 (24.3) | 101 (74.3) | 124 (91.2) | 135 (99.3) | 136 (100) |

Abbreviations: LZ: linezolid; CC: clindamicin; GM: gentamicin; CIP: ciprofloxacin; P: penicillin; E: erythromycin; TE: tetracycline; FOX: cephalosporin;SXT: trimethoprim-sulfamethoxazole. ;

In total, 23 (16.9%) isolates did not produce biofilms, 31 (22.7%) were weak producers, 55 (40.4%) were moderate producers, and 27 (19.8%) were strong producers of biofilms.

We searched for differences between 2 categories of biofilm production (none or weak vs. moderate-strong producers) and antimicrobial susceptibility profiles of the isolates (Table 3). We found no difference in antimicrobial susceptibility among isolates that were moderate or strong biofilm producers.

**Table 3.**
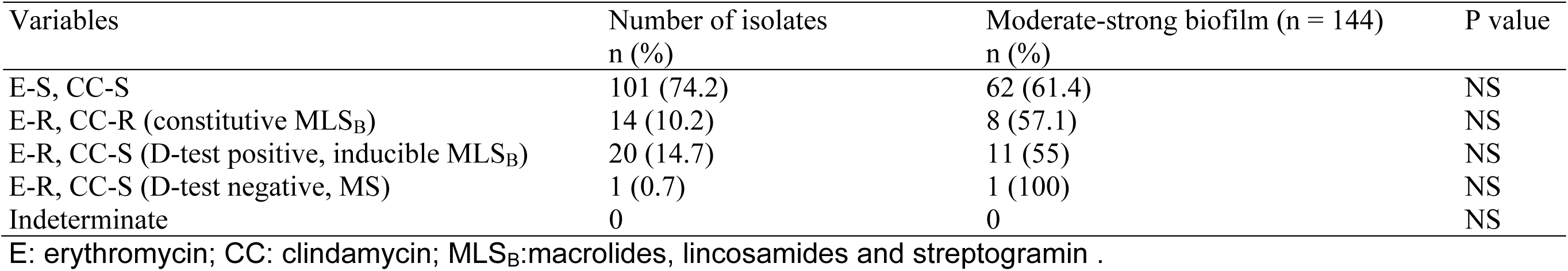
*S. aureus* MLSB sensitivity (n = 243)

| Variables | Number of isolates<br>n (%) | Moderate-strong biofilm (n = 144)<br>n (%) | P value |
| --- | --- | --- | --- |
| E-S, CC-S | 101 (74.2) | 62 (61.4) | NS |
| E-R, CC-R (constitutive MLS<sub>B</sub>) | 14 (10.2) | 8 (57.1) | NS |
| E-R, CC-S (D-test positive, inducible MLS<sub>B</sub>) | 20 (14.7) | 11 (55) | NS |
| E-R, CC-S (D-test negative, MS) | 1 (0.7) | 1 (100) | NS |
| Indeterminate | 0 | 0 | NS |

E: erythromycin; CC: clindamycin; MLS<sub>B</sub>:macrolides, lincosamides and streptogramin .

### Clinical characteristics, antimicrobial susceptibility patterns and biofilm formation

We analyzed the clinical characteristics of patients from whom the isolates were obtained, and the characteristics of the *S. aureus* isolates (Table 4). Patients with non-severe AD at baseline were more likely to have isolates from the initial visit that were resistant to P (p*=*0.02). Those with severe AD at baseline were more likely to have *S. aureus* isolates that exhibited moderate-strong production of biofilms (p*=*0.01).

**Table 4.**
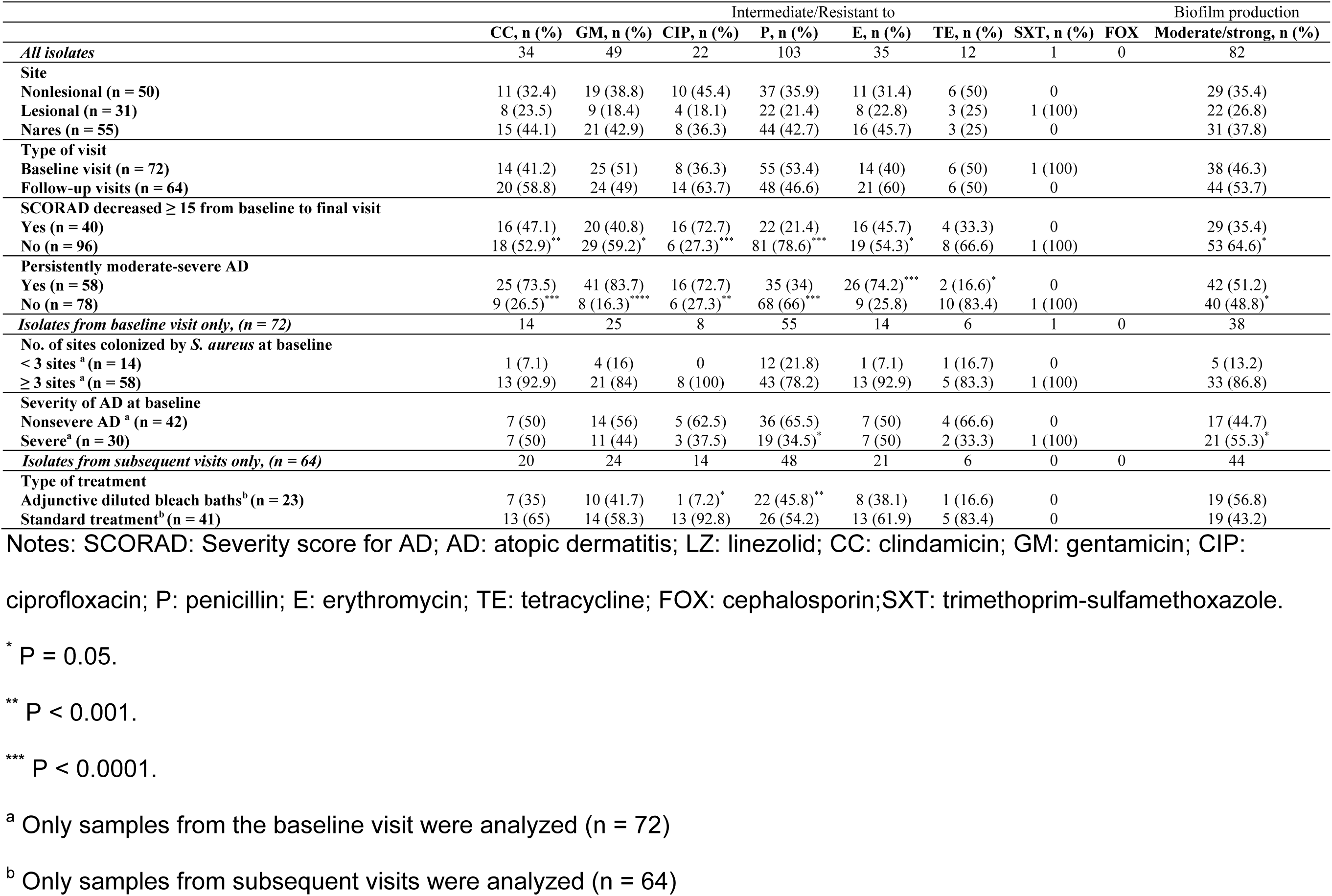
Antibiotic resistance in *S. aureus* isolates and clinical characteristics of the studied population.

Isolates from patients whose SCORAD improved throughout the study were more frequently resistant to P (p=0.001) and more likely to be non or weak producers. (p=0.04). Isolates from those whose SCORAD did not improve accordingly were more frequently resistant to CIP (p<0.0001), GM (p=0.03), CC (p=0.01), and E (p=0.02). These isolates were also more likely to have cMLSB and iMLSB (p<0.0001).

*S. aureus* isolates from patients with persistent moderate–severe AD throughout the study were significantly more frequently resistant (or intermediately resistant) to CIP (p=0.03) but sensitive to P (p<0.0001) and TE (p=0.02). These isolates were also more likely to have cMLSB and iMLSB (p<0.0001) and be moderate-strong biofilm producers (p=0.01). Isolates collected during follow-up visits from patients who did not receive adjunctive treatment with dilute bleach baths were more likely to be resistant to CIP (p<0.0001). These isolates were also more likely to have cMLSB and iMLSB (p=0.01).

Isolates from patients who received adjunctive treatment with dilute bleach baths were more likely to be resistant to P (p=0.004).

## Discussion

In this study of 136 isolates of *S. aureus* from children with AD, we found a predominance of moderate to strong biofilm production capacity (60.2%), which was associated with clinical characteristics such as disease severity at baseline, lack of response to treatment, and persistent moderate–severe AD throughout the study visits. We also found suboptimal rates (<15%) of antimicrobial susceptibility to CC, GM, P, and E.

*S. aureus* is a frequent colonizer in patients with AD; approximately 70% of affected individuals exhibited colonization in lesional skin, compared to 10% of healthy individuals (22, 23). Our understanding of the role of this species in AD has improved recently. The abundance of *S. aureus* is known to be closely associated with flares and loss of microbiome diversity (24), and the bacterium plays a role in epidermal barrier malfunction and inflammation (25, 26).

Moreover, *S. aureus* evolves and adapts within hosts with AD and likely develops additional virulence factors that contribute to adhesion, inflammation and barrier alterations, such as clumping factors and loss of capsule elements (27, 28). Specific clonal complexes have been found in patients with AD and are associated with disease exacerbation (3, 27, 29).

Biofilms produced by *S. aureus* protect it from antimicrobial peptides and phagocytosis, enabling persistence in the host (12). In our study, we found that 60.2% of the isolates were moderate-to-strong biofilm producers, which is similar to previous findings in a study on children with AD (30). One study reported that 67% of *S. aureus* strains were strong biofilm producers, whereas strains from healthy carriers presented lower biofilm production capacity (30). These findings suggest a potential role for biofilms in the pathogenesis of AD (31). Thus, the significant associations between increased biofilm production and disease severity at baseline, lack of response to treatment, and persistent moderate– severe AD throughout the study visits observed herein are relevant. Our results align with those of Di Domenico et al., who reported a correlation between biofilm formation and an increase in the severity of AD lesions. Additionally, they noted that *S. aureus* biofilm production occurred in both the acute and chronic phases of the disease (32). It is hypothesized that biofilms contribute to sweat duct occlusion and skin inflammation by inducing keratinocyte proliferation and cytokine secretion (30). These processes may collectively impair skin barrier function, promote sensitization, and exacerbate pruritus, thereby prolonging the disease (14).

However, biofilm production is also a successful strategy that protects bacteria from environmental danger and treatments such as antibiotics, which potentially contributes to increasing the resistance of *S. aureus* to conventional antibiotics (14) Importantly, all the isolates studied were methicillin-sensitive *S. aureus* (MSSA) strains. However, some studies have reported a low prevalence of MRSA in children with AD (33), along with an absence of multidrug resistance in isolates from both children and adults with AD (34). MRSA strains are associated with more severe infections and exhibit greater complexity in terms of antibiotic resistance. However, it remains controversial whether MRSA strains are inherently more virulent than MSSA strains are (35, 36). We found that more than 50% of the MSSA strains studied produced moderate to strong biofilms. A potential difference in biofilm formation between these two bacterial types has been suggested: a review of the differences in biofilm production between MRSA and MSSA was performed, and among the 20 research articles analyzed, 50% reported that MRSA isolates form biofilms better than MSSA, 45% reported no differences, and only 5% reported that MSSA formed biofilms better than MRSA. There is insufficient evidence to support differences in biofilm formation capacity between MRSA and MSSA isolates. Future analyses should focus on better characterizing the isolates to provide more detailed information on their clonal origin (37).

The ability of strains to form biofilms, combined with their characteristic resistance profile, significantly contributes to an overall increase in bacterial resistance. Biofilms create a protective barrier that impedes the action of antibiotics, and the multidrug resistance profile further amplifies this challenge, severely limiting the availability of therapeutic options (38). Consequently, this synergistic interaction between biofilm formation and multidrug resistance can lead to persistent therapeutic failure, highlighting the urgent need for innovative strategies to combat biofilm-associated infections.

Our findings of suboptimal antimicrobial sensitivity to antibiotics frequently used and recommended to treat AD flares and/or superinfections, such as CC and E, are relevant. The question of whether *S. aureus* in the context of AD has different antimicrobial susceptibility profiles than others has been posed and answered several times. A pivotal study by Allen et al. revealed, similar to our findings, that the antibiotics with the highest percentage of isolates that showed resistance were E (85.7%), CC (80.0%), and levofloxacin (65.7%) (30). However, we obtained no MRSA isolates, unlike studies in other populations, where the rates reached as high as 60% (30). In a recent meta-analysis, the antimicrobial susceptibility of *S. aureus* isolated from patients with AD to commonly prescribed beta-lactams, E, CC, and fusidic acid was found to be <85% (4), a proposed threshold at which an antibiotic should no longer be considered an empirical therapy (39). All of these findings emphasize that antibiotic treatment in patients with AD should be reserved for clinically evident superinfections and always guided by culture.

We observed particular antimicrobial susceptibility profiles for the studied *S. aureus* isolates. In patients who did not have severe AD at baseline, in those whose severity improved throughout the study and in those in whom the disease was not persistently severe, isolates were more frequently resistant to P. Strains of *S. aureus* resistant to P have been documented since 1942 (40), and the current rates of resistance to this antibiotic are between 14% and 26% for MSSA (data from 1997 to 2016) (41), but in other studies, resistance rates as high as 85% have been reported (42, 43). Thus, we expected that the resistance to P of the isolates studied would be high. Resistance to P does not necessarily imply increased virulence of *S. aureus,* but in some reports, the coexistence of P resistance in MSSA isolates and the presence of Panton-Valentine leukocidin has been reported (44, 45). Although this agent is seldom used to treat patients with AD and SSTIs, we decided to include it in testing, expecting a low frequency of MRSA isolates in the cohort of patients studied. Characterizing antimicrobial susceptibility to P offers relevant epidemiological information, indicating that the susceptibility profiles of AD patients to beta-lactams are different from those of patients with other types of infection (41, 46).

Patients who did not exhibit improvement harbored isolates more likely resistant to CIP, GM, CC, and E. High rates of E and CC resistance have been observed for MSSA, but a relationship with microbiological treatment has not been established. (42, 43, 46). Further studies could be useful in confirming or ruling out this association.

Data on MLSB phenotypes in *S. aureus* from AD patients are limited. In this collection, the iMLSB phenotype was observed in 14.7% of the isolates, and notably, all the isolates were MSSA. The distribution of the iMLSB phenotype is reported to differ between MRSA and MSSA (18); in MRSA, the frequency varies from 0 to 76.4% (47–49), whereas in MSSA, it ranges from 5.9 to 66% (48, 50, 51).

In China, Japan and Brazil, the *erm*A and *erm*B genes are predominant among MRSA and MSSA (48, 50, 52); however, in studies on isolates from other regions of Asia and Africa, *erm*C was the most frequently detected gene (40.7-72.4%) (47, 53, 54). Information from Mexico about the genes involved in MLSB phenotypes is limited; in a collection of 21 MRSA isolates obtained from catheter-associated infections, the coexistence of *erm*A and *erm*B was observed (55). In this study, the *erm*A gene was detected in 80% of the isolates with the iMLSB phenotype. No gene related to the iMLSB phenotype was amplified from four isolates, which indicates that another allele of *erm* could be involved, such as *emr*T, which has been detected in isolates from the hospital environment, nostrils of inpatients and health workers (54, 56).

We found a significant association between the cMLSB and iMLSB phenotypes in isolates from patients who showed no improvement throughout the study, who had persistent moderate–severe AD, and who did not receive diluted bleach baths. These findings merit further study in prospective studies with larger sample sizes.

All of these findings support the need to develop specific targeted treatments with different mechanisms of action than those of antibiotics to decrease the abundance of *S. aureus* and its virulence and pathogenic factor levels, particularly those related to biofilm production and quorum sensing.

Sodium hypochlorite, which is commonly used in the management of AD at concentrations ranging from 0.005-0.006% as an adjunctive treatment, has recently been shown in a meta-analysis to be effective in decreasing AD severity as well as *S. aureus* abundance (57). It has also proven effective against *S. aureus* biofilms. However, the concentrations required to eliminate *S. aureus* biofilms are greater than those for planktonic cells at concentrations between 0.01% and 0.08%. *In vitro* eradication of *S. aureus* biofilms was achieved at concentrations ranging from 0.01% to 0.16%, much higher than those currently used in bleach baths for patients with AD (0.006%) (58, 59). We found that adjunctive treatment with dilute bleach baths at follow-up visits was associated with resistance to CIP and with constitutive and inducible MLSB. Although there are no studies supporting this association in *S. aureus,* previous studies on *Candida albicans* have reported that adaptation to sodium hypochlorite renders it ineffective (60). We believe our findings are relevant and warrant further study to establish whether sodium hypochlorite may play a potential role in gene expression, adaptation, or other mechanisms of resistance in *S. aureus*.

In terms of limitations, our results might be skewed by selection bias, as all the isolates were obtained from patients cared for at a tertiary institution who might have particular antimicrobial susceptibility profiles and biofilm production phenotypes. Additionally, we did not test for other relevant antibiotics or perform functional studies that might have shed more light on *S. aureus* pathogenicity and virulence in the context of AD.

However, data on the specific characteristics of *S. aureus* as a colonizer and pathogen in the skin of patients with AD are scarce. In Mexico, data on the antimicrobial susceptibility patterns of the bacteria causing soft tissue and skin infections are scarce, even more so for bacteria from the skin of patients with AD. Our findings contribute to epidemiological data on antimicrobial susceptibility profiles, are essential for establishing regional treatment guidelines, and offer further insights into the role *S. aureus* in the pathophysiology of AD.

## Conclusions

In conclusion, 60.2% of *S. aureus* isolates from children with AD were moderate–strong biofilm producers. Furthermore, moderate-strong biofilm production was significantly associated with severe AD at baseline, a lack of response to treatment, and persistent moderate-severe AD throughout the study visits. We also found suboptimal rates of resistance to frequently used and recommended antibiotics in the context of AD, such as CC, P and E, which might be a consequence of the successful survival strategies offered by biofilm production to the bacteria. Our findings indicate that adequate treatment with SSTIs in patients with AD should always be guided by culture and antibiograms. Moreover, these results highlight the need to develop targeted therapies to treat colonization and superinfections in patients with AD while not promoting antibiotic resistance, such as specifically targeting biofilm production and quorum sensing as key strategies to manage colonization and treatment of recurrent infections in AD. Further research on host‒ microbial interactions and their implications for AD and increasing our understanding of staphylococcal biofilms in the context of AD will allow the development of targeted treatments to reduce skin colonization, improve barrier function, weaken the immune response and reduce flares while not promoting antibiotic resistance.

## Acknowledgments

We acknowledge support the Mexican Government Ministry of Taxes Program E022 for Health Research Development (to M.T.G.-R.).

## Author contribution

C.R-G. cultured bacteria from clinical samples and extracted DNA and designed the biofilm assay. A. A-A designed and performed all the gene determination by PCR. A. P-C. and A. B-O performed the antimicrobial susceptibility testing. D. CH-C performed the biofilm assay. M.T.G.-R. designed the clinical cohort, enrolled patients and collected clinical samples, Funding Acquisition and wrote the manuscript with feedback from C.R-G. and A. A-A.

## References

1. Tollefson MM, Bruckner AL, Cohen BA, Antaya R, Bruckner A, Horii K, Silverberg NB, Wright T. 2014. Atopic dermatitis: skin-directed management. Pediatrics 134:e1735–e1744.

2. Kong HH, Oh J, Deming C, Conlan S, Grice EA, Beatson MA, Nomicos E, Polley EC, Komarow HD, Nisc Comparative Sequence Program, Murray PR, Turner ML, Segre JA. 2012. Temporal shifts in the skin microbiome associated with disease flares and treatment in children with atopic dermatitis. Genome Res 22:850–859.

3. Paller AS, Kong HH, Seed P, Naik S, Scharschmidt TC, Gallo RL, Luger T, Irvine AD. 2019. The microbiome in patients with atopic dermatitis. J Allergy Clin Immunol 143:26–35.

4. Elizalde-Jiménez IG, Ruiz-Hernández FG, Carmona-Cruz SA, Pastrana-Arellano E, Aquino-Andrade A, Romo-González C, Arias-de la Garza E, Álvarez-Villalobos NA, García-Romero MT. 2024. Global antimicrobial susceptibility patterns of staphylococcus aureus in atopic dermatitis: a systematic review and meta-analysis. JAMA Dermatol 160:1171–1181.

5. Dukic VM, Lauderdale DS, Wilder J, Daum RS, David MZ. 2013. Epidemics of community-associated methicillin-resistant Staphylococcus aureus in the United States: a meta-analysis. PLoS One 8:e52722.

6. Jung MY, Chung JY, Lee HY, Park J, Lee DY, Yang JM. 2015. Antibiotic susceptibility of staphylococcus aureus in atopic dermatitis: current prevalence of methicillin-resistant staphylococcus aureus in korea and treatment strategies. Ann Dermatol 27:398–403.

7. Edslev SM, Clausen ML, Agner T, Stegger M, Andersen PS. 2018. Genomic analysis reveals different mechanisms of fusidic acid resistance in Staphylococcus aureus from Danish atopic dermatitis patients. J Antimicrob Chemother 73:856– 861.

8. Bessa GR, Quinto VP, Machado DC, Lipnharski C, Weber MB, Bonamigo RR, D’Azevedo PA. 2016. Staphylococcus aureus resistance to topical antimicrobials in atopic dermatitis. An Bras Dermatol 91:604–610.

9. Alzolibani A, Al Robaee A, Al Shobaili H, Bilal J, Ahmad M, Saif GB. 2012. Documentation of vancomycin-resistant Staphylococcus aureus (VRSA) among children with atopic dermatitis in the Qassim region, Saudi Arabia. Acta Dermatovenerol Alp Pannonica Adriat 21:51–53.

10. Velázquez-Meza ME, Mendoza-Olazarán S, Echániz-Aviles G, Camacho-Ortiz A, Martínez-Reséndez MF, Valero-Moreno V, Garza-González E. 2017. Chlorhexidine whole-body washing of patients reduces methicillin-resistant Staphylococcus aureus and has a direct effect on the distribution of the ST5-MRSA-II (New York/Japan) clone. J Med Microbiol 66:721–728.

11. Gonzalez T, Stevens ML, Baatyrbek Kyzy A, Alarcon R, He H, Kroner JW, Spagna D, Grashel B, Sidler E, Martin LJ, Biagini Myers JM, Khurana Hershey GK, Herr AB. 2021. Biofilm propensity of Staphylococcus aureus skin isolates is associated with increased atopic dermatitis severity and barrier dysfunction in the MPAACH pediatric cohort. Allergy 76:302–313.

12. Sonesson A, Przybyszewska K, Eriksson S, Mörgelin M, Kjellström S, Davies J, Potempa J, Schmidtchen A. 2017. Identification of bacterial biofilm and the Staphylococcus aureus derived protease, staphopain, on the skin surface of patients with atopic dermatitis. Sci Rep 7:8689.

13. Peng Q, Tang X, Dong W, Sun N, Yuan W. 2022. A review of biofilm formation of staphylococcus aureus and its regulation mechanism. Antibiotics (Basel, Switzerland) 12:12.

14. Gonzalez T, Myers JMB, Herr AB, Hershey GKK. 2017. Staphylococcal biofilms in atopic dermatitis. Curr Allergy Asthma Rep 17:81.

15. Chopra R, Vakharia PP, Sacotte R, Patel N, Immaneni S, White T, Kantor R, Hsu DY, Silverberg JI. 2017. Severity strata for eczema area and severity index (EASI), modified EASI, scoring atopic dermatitis (SCORAD), objective SCORAD, atopic dermatitis severity index and body surface area in adolescents and adults with atopic dermatitis. Br J Dermatol 177:1316–1321.

16. CLSI. Performance Standards for Antimicrobial Susceptibility Testing. 33th ed. CLSI supplement M100. Clinical and Laboratory Standards Institute; 2023.

17. CLSI. Performance Standards for Antimicrobial Susceptibility Testing. 29th ed. CLSI supplement M100. Clinical and Laboratory Standards Institute; 2019.

18. Miklasińska-Majdanik M. 2021. Mechanisms of resistance to macrolide antibiotics among *Staphylococcus aureus*. Antibiotics (Basel, Switzerland) 10:1406.

19. Fri J, Njom HA, Ateba CN, Ndip RN. 2020. Antibiotic resistance and virulence gene characteristics of methicillin-resistant *Staphylococcus aureus* (MRSA) isolated from healthy edible marine fish. Int J Microbiol 2020:9803903.

20. Khodabandeh M, Mohammadi M, Abdolsalehi MR, Alvandimanesh A, Gholami M, Bibalan MH, Pournajaf A, Kafshgari R, Rajabnia R. 2019. Analysis of resistance to macrolide-lincosamide-streptogramin B among mecA-positive *Staphylococcus aureus* isolates. Osong Public Health Res Perspect 10:25–31.

21. Singh AK, Prakash P, Achra A, Singh GP, Das A, Singh RK. 2017. Standardization and Classification of In vitro Biofilm Formation by Clinical Isolates of *Staphylococcus aureus*. J Glob Infect Dis 9:93–101.

22. Harkins CP, Holden MTG, Irvine AD. 2019. Antimicrobial resistance in atopic dermatitis. Ann Allergy Asthma Immunol 122:236–240.

23. Rangel SM, Paller AS. 2018. Bacterial colonization, overgrowth, and superinfection in atopic dermatitis. Clin Dermatol 36:641–647.

24. Khadka VD, Key FM, Romo-González C, Martínez-Gayosso A, Campos-Cabrera BL, Gerónimo-Gallegos A, Lynn TC, Durán-McKinster C, Coria-Jiménez R, Lieberman TD, García-Romero MT. 2021. The skin microbiome of patients with atopic dermatitis normalizes gradually during treatment. Front Cell Infect Microbiol 11:720674.

25. Hwang J, Thompson A, Jaros J, Blackcloud P, Hsiao J, Shi VY. 2021. Updated understanding of *Staphylococcus aureus* in atopic dermatitis: from virulence factors to commensals and clonal complexes. Exp Dermatol 30:1532–1545.

26. Simpson EL, Villarreal M, Jepson B, Rafaels N, David G, Hanifin J, Taylor P, Boguniewicz M, Yoshida T, De Benedetto A, Barnes KC, Leung DYM, Beck LA. 2018. Patients with atopic dermatitis colonized with staphylococcus aureus have a distinct phenotype and endotype. J Investig Dermatol 138:2224–2233.

27. Fleury OM, McAleer MA, Feuillie C, Formosa-Dague C, Sansevere E, Bennett DE, Towell AM, McLean WHI, Kezic S, Robinson DA, Fallon PG, Foster TJ, Dufrêne YF, Irvine AD, Geoghegan JA. 2017. Clumping factor B promotes adherence of staphylococcus aureus to corneocytes in atopic dermatitis. Infect immun 85:e00994–16.

28. Key FM, Khadka VD, Romo-González C, Blake KJ, Deng L, Lynn TC, Lee JC, Chiu IM, García-Romero MT, Lieberman TD. 2023. On-person adaptive evolution of Staphylococcus aureus during treatment for atopic dermatitis. Cell Host Microbe 31:593–603.e597.

29. Clausen ML, Edslev SM, Andersen PS, Clemmensen K, Krogfelt KA, Agner T. 2017. *Staphylococcus aureus* colonization in atopic eczema and its association with filaggrin gene mutations. Br J Dermatol 177:1394–1400.

30. Allen HB, Vaze ND, Choi C, Hailu T, Tulbert BH, Cusack CA, Joshi SG. 2014. The presence and impact of biofilm-producing staphylococci in atopic dermatitis. JAMA Dermatol 150:260–265.

31. Conte AL, Brunetti F, Marazzato M, Longhi C, Maurizi L, Raponi G, Palamara AT, Grassi S, Conte MP. 2023. Atopic dermatitis-derived Staphylococcus aureus strains: what makes them special in the interplay with the host. Front Cell Infect Microbiol 13:1194254.

32. Di Domenico EG, Cavallo I, Bordignon V, Prignano G, Sperduti I, Gurtner A, Trento E, Toma L, Pimpinelli F, Capitanio B, Ensoli F. 2018. Inflammatory cytokines and biofilm production sustain *Staphylococcus aureus* outgrowth and persistence: a pivotal interplay in the pathogenesis of atopic dermatitis. Sci Rep 8:9573.

33. Rezaei M, Chavoshzadeh Z, Haroni N, Armin S, Navidinia M, Mansouri M, Shamshiri AR, Mesdaghi M, Eshgh FA. 2013. Colonization with methicillin resistant and methicillin sensitive *Staphylococcus aureus* subtypes in patients with atopic dermatitis and its relationship with severity of eczema. Arch Pediatr Infect Dis 1:53–56.

34. Ogonowska P, Szymczak K, Empel J, Urbaś M, Woźniak-Pawlikowska A, Barańska-Rybak W, Świetlik D, Nakonieczna J. 2023. Staphylococcus aureus from atopic dermatitis patients: its genetic structure and susceptibility to phototreatment. Microbiol Spectr 11:e0459822.

35. Fooladi AAI, Ashrafi E, Tazandareh SG, Koosha RZ, Rad HS, Amin M, Soori M, Larki RA, Choopani A, Hosseini HM. 2015. The distribution of pathogenic and toxigenic genes among MRSA and MSSA clinical isolates. Microb Pathog 81:60– 66.

36. Rozgonyi F, Kocsis E, Kristóf K, Nagy K. 2007. Is MRSA more virulent than MSSA? Clin Microbiol Infect 13:843–845.

37. Hernández-Cuellar E, Tsuchiya K, Valle-Ríos R, Medina-Contreras O. 2023. Differences in biofilm formation by methicillin-resistant and methicillin-susceptible *Staphylococcus aureus* strains. Diseases (Basel, Switzerland) 11:160.

38. Silva V, Hermenegildo S, Ferreira C, Manaia CM, Capita R, Alonso-Calleja C, Carvalho I, Pereira JE, Maltez L, Capelo JL, Igrejas G, Poeta P. 2020. Genetic characterization of methicillin-resistant *Staphylococcus aureus* isolates from human bloodstream infections: detection of MLS(B) resistance. Antibiotics (Basel, Switzerland) 9:375.

39. Cressman AM, MacFadden DR, Verma AA, Razak F, Daneman N. 2018. Empiric antibiotic treatment thresholds for serious bacterial infections: a scenario-based survey study. Clin Infect Dis 69:930–937.

40. Lade H, Kim JS. 2023. Molecular determinants of β-lactam resistance in methicillin-resistant *Staphylococcus aureus* (MRSA): an updated review. Antibiotics (Basel, Switzerland) 12:1362.

41. Diekema DJ, Pfaller MA, Shortridge D, Zervos M, Jones RN. 2019. Twenty-year trends in antimicrobial susceptibilities among staphylococcus aureus from the SENTRY antimicrobial surveillance program. Open Forum Infect Dis 6:S47–S53.

42. Dai C, Ji W, Zhang Y, Huang W, Wang H, Wang X. 2024. Molecular characteristics, risk factors, and clinical outcomes of methicillin-resistant Staphylococcus aureus infections among critically ill pediatric patients in Shanghai, 2016-2021. Front Pediatr 12:1457645.

43. McNeil JC, Sommer LM, Joseph M, Hulten KG, Kaplan SL. 2024. Penicillin susceptibility among Staphylococcus aureus skin and soft tissue infections at a children’s hospital. Microbiol Spectr 12:e0086924.

44. Darboe S, Dobreniecki S, Jarju S, Jallow M, Mohammed NI, Wathuo M, Ceesay B, Tweed S, Basu Roy R, Okomo U, Kwambana-Adams B, Antonio M, Bradbury RS, de Silva TI, Forrest K, Roca A, Lawal BJ, Nwakanma D, Secka O. 2019. Prevalence of panton-valentine leukocidin (PVL) and antimicrobial resistance in community-acquired clinical *Staphylococcus aureus* in an urban gambian hospital: a 11-year period retrospective pilot study. Front Cell Infect Microbiol 9:170.

45. Kilani AM, Alabi ED, Adeleke OE. 2024. Coexistence of the blaZ gene and selected virulence determinants in multidrug-resistant Staphylococcus aureus: insights from three Nigerian tertiary hospitals. BMC Infect Dis 24:1269.

46. de Menezes IL, Pone SM, Pone MVDS. 2024. Clinical, demographic characteristics and antimicrobial resistance profile of Staphylococcus aureus isolated in clinical samples from pediatric patients in a tertiary hospital in Rio de Janeiro: 7-year longitudinal study. BMC Infect Dis 24:1081–1081.

47. Assefa M. 2022. Inducible clindamycin-resistant *Staphylococcus aureus* strains in Africa: a systematic review. Int J Microbiol 2022:1835603.

48. Pereira JNDP, Rabelo MA, Lima JLDC, Neto AMB, Lopes ACDS, Maciel MAV. 2016. Phenotypic and molecular characterization of resistance to macrolides, lincosamides and type B streptogramin of clinical isolates of Staphylococcus spp. of a university hospital in Recife, Pernambuco, Brazil. Braz J Infect Dis 20:276– 281.

49. Timsina R, Shrestha U, Singh A, Timalsina B. 2020. Inducible clindamycin resistance and erm genes in Staphylococcus aureus in school children in Kathmandu, Nepal. Future Sci 7:FSO361.

50. Wang L, Liu Y, Yang Y, Huang G, Wang C, Deng L, Zheng Y, Fu Z, Li C, Shang Y, Zhao C, Sun M, Li X, Yu S, Yao K, Shen X. 2012. Multidrug-resistant clones of community-associated meticillin-resistant Staphylococcus aureus isolated from Chinese children and the resistance genes to clindamycin and mupirocin. J Med Microbiol 61:1240–1247.

51. Elkammoshi AM, Ghasemzadeh-Moghaddam H, Nordin SA, Taib NM, Subbiah SK, Neela V, Hamat RA. 2016. A low prevalence of inducible macrolide, lincosamide, and streptogramin B resistance phenotype among methicillin-susceptible *Staphylococcus aureus* isolated from Malaysian patients and healthy individuals. Jundishapur J Microbiol 9:e37148.

52. Nahar L, Hagiya H, Nada T, Iio K, Gotoh K, Matsushita O, Otsuka F. 2023. Prevalence of inducible macrolide, lincosamide, and streptogramin B (inducible MLSB) resistance in clindamycin-susceptible *Staphylococcus aureus* at Okayama university hospital. Acta Med Okayama 77:1–9.

53. Goudarzi M, Tayebi Z, Fazeli M, Miri M, Nasiri MJ. 2020. Molecular characterization, drug resistance and virulence analysis of constitutive and inducible clindamycin resistance *Staphylococcus aureus* strains recovered from clinical samples, Tehran - iran. Infect Drug Resist 13:1155–1162.

54. Manouchehrifar M, Khademi F, Doghaheh HP, Habibzadeh S, Arzanlou M. 2023. Macrolide-lincosamide resistance and virulence genes in *Staphylococcus aureus* isolated from clinical specimens in Ardabil, Iran. Iran J Pathol 18:415–424.

55. Paniagua-Contreras GL, Monroy-Pérez E, Vaca-Paniagua F, Rodríguez-Moctezuma JR, Negrete-Abascal E, Vaca S. 2014. Implementation of a novel in vitro model of infection of reconstituted human epithelium for expression of virulence genes in methicillin-resistant Staphylococcus aureus strains isolated from catheter-related infections in Mexico. Ann Clin Microbiol Antimicrob 13:6.

56. Laceb ZM, Diene SM, Lalaoui R, Kihal M, Chergui FH, Rolain JM, Hadjadj L. 2022. Genetic diversity and virulence profile of methicillin and inducible clindamycin-resistant *Staphylococcus aureus* isolates in Western Algeria. Antibiotics (Basel, Switzerland) 11:971.

57. Bakaa L, Pernica JM, Couban RJ, Tackett KJ, Burkhart CN, Leins L, Smart J, Garcia-Romero MT, Elizalde-Jiménez IG, Herd M, Asiniwasis RN, Boguniewicz M, De Benedetto A, Chen L, Ellison K, Frazier W, Greenhawt M, Huynh J, LeBovidge J, Lind ML, Lio P, O’Brien M, Ong PY, Silverberg JI, Spergel JM, Wang J, Begolka WS, Schneider L, Chu DK. 2022. Bleach baths for atopic dermatitis. Ann Allergy Asthma Immunol 128:660–668.e669.

58. Eriksson S, van der Plas MJA, Mörgelin M, Sonesson A. 2017. Antibacterial and antibiofilm effects of sodium hypochlorite against *Staphylococcus aureus* isolates derived from patients with atopic dermatitis. Br J Dermatol 177:513–521.

59. Wong SM, Ng TG, Baba R. 2013. Efficacy and safety of sodium hypochlorite (bleach) baths in patients with moderate to severe atopic dermatitis in Malaysia. J Dermatol 40:874–880.

60. Karpiński TM, Korbecka-Paczkowska M, Ożarowski M, Włodkowic D, Wyganowska ML, Seremak-Mrozikiewicz A, Cielecka-Piontek J. 2024. Adaptation to sodium hypochlorite and potassium permanganate may lead to their ineffectiveness against candida albicans. Pharmaceuticals (Basel, Switz

